# HiMoRNA and RNA-Chrom integration: Chromatin-Associated LncRNAs in Genome-Wide Epigenetic Regulation

**DOI:** 10.1101/2024.05.02.592208

**Authors:** Ivan S. Ilnitskiy, Grigory K. Ryabykh, Daria A. Marakulina, Andrey A. Mironov, Yulia A. Medvedeva

## Abstract

Long non-coding RNAs (lncRNAs) significantly contribute to genome structure and regulation. Many lncRNAs are known to interact with chromatin and in this way to affect gene expression patterns through epigenetic regulation. Still, experimental protocols for lncRNA-chromatin interactions do not provide any insight into the mechanisms of lncRNA-based genome-wide regulation. Here we present an integration of HiMoRNA – a resource containing correlated lncRNA-epigenetic changes in specific genomic locations genome-wide, – and RNA-Chrom, a resource featuring uniformly processed experimental data on RNA-chromatin interactions. Our integration approach allows generating interpretable and experimentally supported hypotheses on the mechanisms of lncRNA epigenetic regulation of gene expression. For this integration we have tailored the interface of HiMoRNA such that for many lncRNAs experimentally detected RNA-chromatin contacts are available from RNA-Chrom for browsing, analysis and downloading. HiMoRNA peaks supported by RNA-Chrom contacts can be explained by external experimental data. We believe that the integration of HiMoRNA and RNA-Chrom is a convenient and valuable approach that can provide experimental and mechanistic insights and greatly facilitate functional annotation of lncRNAs.

## 1. Introduction

A considerable number of human cells transcribe a wide variety of long noncoding RNAs (lncRNAs), comparable in number to that of protein-coding genes [1]. Due to their low expression, tissue-specificity and evolutionary conservation, lncRNAs are challenging to functionally annotate [2,3]. Nevertheless, other characteristics of lncRNAs are often conserved, including synteny with neighbouring genes, similarity of short sequence fragments, and secondary structure [4]. These observations suggest that lncRNAs could potentially be functional. Additionally, indirect evidence indicates that lncRNAs are tightly regulated and participate in diverse molecular mechanisms.

The majority of lncRNAs has been detected to interact with chromatin and are essential for the epigenetic control of specific genomic loci as well as the organisation of the chromosomes [5–8]. The identification of functional genomic targets for chromatin-interacting lncRNAs is currently critical. Previously, we developed HiMoRNA [9], a multi-omics resource integrating correlated lncRNA–epigenetic changes in specific genomic locations genome-wide to computationally predict the effects of lncRNA on epigenetic modifications and gene expression. However, to generate reasonable hypotheses for further experimental validation, a list of 5 million lncRNA-correlated HiMoRNA peaks has to be narrowed down to detect the most reliable ones. Experimental methods for the detection of RNA-chromatin interactions present a valuable data source for this purpose.

A number of experimental methods have been developed to identify ncRNAs contacting chromatin and their target regions. For simplicity, we can divide them into two groups: “one-to-all” [8–14] and “all-to-all” [15–20]. The first group of methods determines the contacts of a specific RNA with chromatin, while the second aims to uncover all possible RNA-DNA contacts in a cell [21]. Experimentally detected lncRNA–chromatin interactions have been uploaded into RNA-Chrom DB [22], a web-based resource that provides a variety of information on the RNA-chromatin interactions and two types of the data analysis (“from RNA” and “from DNA”), which can be used for research purposes. It is regrettable that all of these experimental methods suffer from a high level of false-positives and do not uncover mechanisms of lncRNA-associated epigenetic regulation. Furthermore, “all-to-all” methods frequently fail to detect contacts for rarely expressed RNAs and are overloaded with the contacts of nascent transcripts. In order to compensate for the issues of HiMoRNA and RNA-Chrom and to improve the characterization of lncRNA in epigenetic regulation, we have integrated these databases. Our approach enables the generation of interpretable hypotheses regarding the mechanisms of lncRNA epigenetic regulation of gene expression, supported by experimental data on RNA-chromatin interactions. For this integration effort, we have modified the interface of HiMoRNA in a way that for 4124 out of 4145 HiMoRNA lncRNAs, there is now access to experimentally supported RNA-chromatin contacts in RNA-Chrom for browsing, analysis and downloading. The results of the use-case analysis indicate that the integration of HiMoRNA and RNA-Chrom represents a convenient and valuable resource that can provide experimental and mechanistic insights into lncRNA functions and greatly facilitate their functional annotation. The HiMoRNA database is available to users at https://himorna.fbras.ru (accessed on 1 April 2024).

## 2. Materials and Methods

### 2.1. Gene names correspondence

The HiMoRNA and RNA-Chrom databases utilize disparate versions of the GENCODE gene annotations, namely gencode.v31.basic.annotation.gtf and gencode.v35.basic.annot6a1tion.gtf, respectively. To ascertain the correspondence between genes across different annotations, we employed three distinct approaches: (1) same gene names; (2) same gene IDs; and (3) overlap of coordinates between two genes. Unfortunately, gene names and coordinates in different annotation sources do not always match. To address this issue, we intersected 4145 lncRNA genes from HiMoRNA with 60619 genes from RNA-Chrom by genomic coordinates using bedtools (intersect command), resulting in 6778 gene pairs. Two genes from the HiMoRNA did not pass this stage (ENSG00000267034.1, ENSG00000280076.1). Subsequently, for each gene pair from different annotation sources, the Jaccard index was calculated as a ratio of the length of the genes’ overlap to the length of their union. Using the Jaccard index threshold of 0.99, we identified 4100 unique matches (Supplementary Figure 1). Of the remaining 47 HiMoRNA genes, we were able to unambiguously match 24 genes with genes from RNA-Chrom by gene_name. In total, we obtained 4124 lncRNA in common in both HiMoRNA and RNA-Chrom DB. The gene correspondence table (lncRNA correspondence table) is available for download from the HiMoRNA web-based resource (Supplementary Table 1).

### 2.2. EZH2-associated chromatin interactions

To facilitate interpretability of the RNA-chromatin contacts we used Red-ChIP data [23] (available in the GEO repository under accession GSE174474, samples GSM5315228 and GSM5315229). The Red-ChIP method captures the RNA-chromatin contacts mediated by a specific protein, in our case, EZH2, a component of the PRC2 complex that establishes H3K27me3. The primary processing of the dataset was made following the established pipeline for RNA-Chrom data. To increase the reliability of the EZH2-mediated contacts of lncRNA PVT1 we performed peak-calling using the BaRDIC program (version cited as per preprint DOI [24]). The analysis parameters were set to use *--qval_type all* and a threshold of 1 for the q-value (*--qval_threshold 1*). In this way we detected 3242 PVT1 peaks, indicating potential regions of interest within the chromatin landscape influenced by EZH2.

## 3. Results

### 3.1. Databases integration

Given that HiMoRNA provides millions of peaks, it is reasonable to focus the downstream analysis on the most reliable ones. To facilitate this, we integrated HiMoRNA peaks with the RNA-chromatin contacts available from RNA-Chrom. To do so, we established a one-to-one correspondence between genes from HiMoRNA DB and RNA-Chrom DB (see the Materials and Methods, “Gene names correspondence”). With this approach, HiMoRNA generates a request to the RNA-Chrom DB for a given RNA with URL (e.g. https://rnachrom2.bioinf.fbb.msu.ru/basic_graphical_summary_dna_filter?locus=chrX:23456-9624253566&name=XIST&rnaID=227896&organism=Homo+sapiens) containing the necessary parameters (locus, name, rnaID, organism). RNA-Chrom is now able to process such URLs and to provide the results in a new window. Overall integration schema is presented at Figure 1.

**Figure 1.**
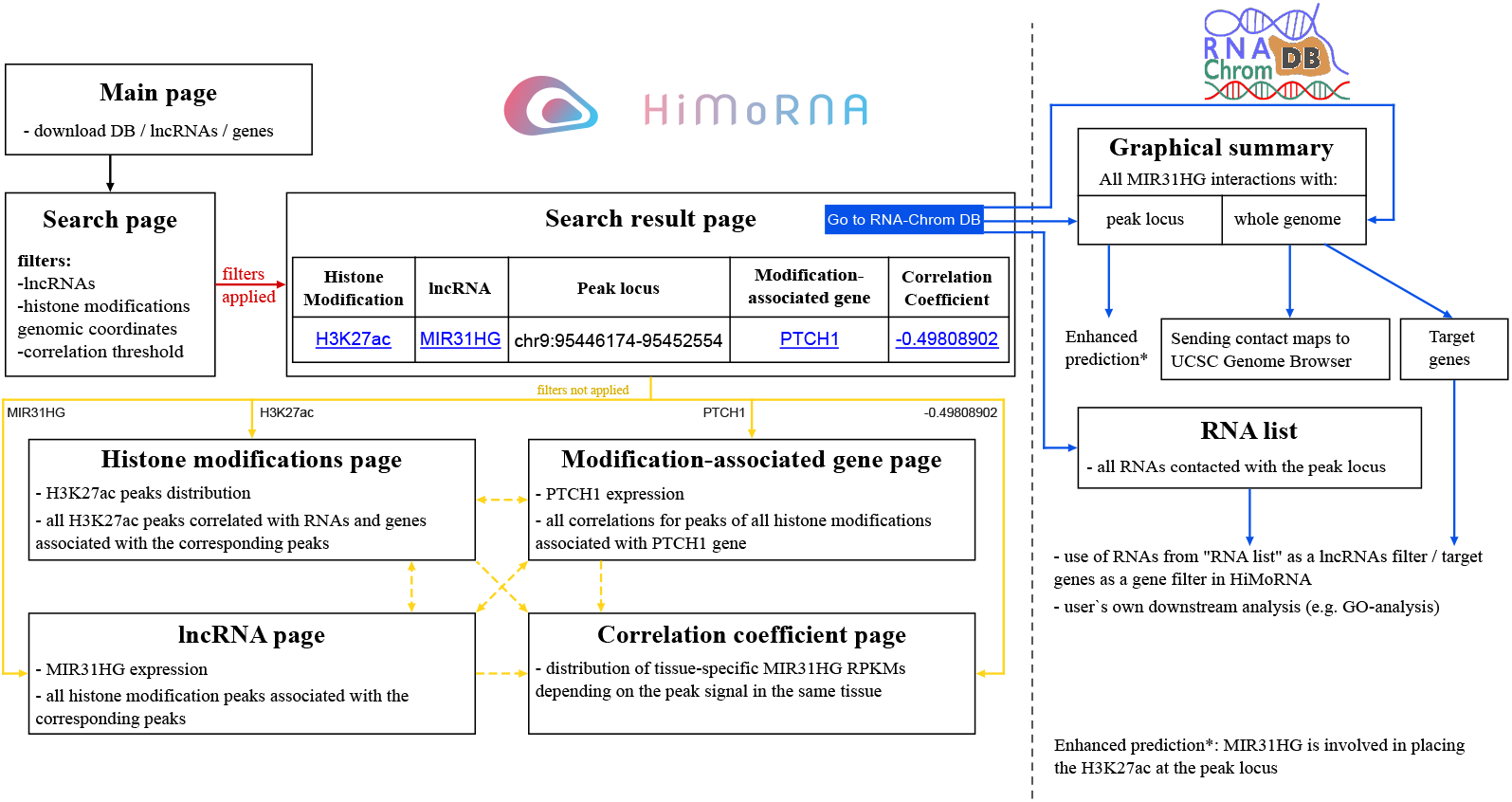
Integrated HiMoRNA – RNA-Chrom databases usage scenario. Rectangles represent web pages, arrows represent transitions between them.

For the purpose of ensuring smooth integration we have improved HiMoRNA’s interface and the efficiency. Now the user can download the “Gene table” and the “lncRNA correspondence table”. The user can search for genes/lncRNAs of interest by genomic coordinates in these files, we provided this option since the user’s Ensembl ID or lncRNA name (as well as gene IDs or names) may not match those in HiMoRNA.

### 3.2. Use case

The primary objective of integrating HiMoRNA and RNA-Chrom is to validate the potential role of an lncRNA in epigenetic regulation by examining its physical interaction with a genomic region in proximity to the correlated peak of a particular histone modification. The following section presents case studies that have been corroborated by external experimental data, which substantiates the regulatory role of a particular lncRNA in histone modification.

#### 3.2.1. MIR31HG lncRNA

The long non-coding RNA MIR31HG is a known regulator of the histone marks H3K1me1, H3K4me3 and H3K27ac. Reduced levels of H3K4me1 and H3K27ac in the enhancer region of the GLI2 gene, as well as of H3K4me3 and H3K27ac in the promoter region of the FABP4 gene have been reported after the MIR31HG knockdown [25,26]. This observation can be verified by using our integration of HiMoRNA with RNA-Chrom. To do so, a query for the lncRNA MIR31HG is created, histone marks H3K4me1 and H3K27ac are selected, and the coordinates of the target genes are specified with an upstream region extended by 10 Kb in the field of genomic coordinates (Supplementary Figure 2A). As a result, the HiMoRNA database generates a table displaying H3K27ac and H3K4me1 peaks correlated with MIR31HG expression in different tissues (Supplementary Figure 2B). We observed that a highly correlated peak is present in the upstream region of GLI2 for each mark. A specific peak is then chosen by clicking the radio button to the left of the peak, and utilising the “Go to RNA-Chrom DB” button (Supplementary Figure 2C) enables the user to verify experimentally detected MIR31HG contacts with the chromatin in the locus of this peak. The user can find all target genes in the vicinity of the selected experiment by selecting one of them in the top table and clicking on the “All target genes” button on the right (Supplementary Figure 2D). The resulting table depicts the interactions of MIR31HG with chromatin in the gene body of the GLI2 gene (Supplementary Figure 2E). A step-by-step analysis is provided in Supplementary Table 2.

To ascertain whether the integration of HiMoRNA and RNA-Chrom can yield novel and valuable biological insights, we assumed that MIR31HG might regulate not only GLI2 but also other genes belonging to the Sonic hedgehog pathway, a pathway in which GLI2 is also involved. To this end, we identified genes of interest using the KEGG Orthology database [27] and constructed a new query for MIR31HG and H3K4me1 and H3K27ac histone marks, including the names of genes or their genomic coordinates in the corresponding fields. Consequently, from HiMoRNA, we obtained a table of 162 entries, which can be confirmed using the RNA-Chrom resource. For instance, one can examine the chromatin contacts of MIR31HG in the locus of H3K27ac peak_963553 (chr9:95446174-95452554) in the PTCH1 gene, this gene encodes a receptor for Sonic hedgehog. Therefore, the integration can be successfully used to find predictions of lncRNAs acting as regulators of histone modification that require experimental validation.

#### 3.2.2. PVT1 lncRNA

The lncRNA PVT1 inhibits the expression of the large tumor suppressor kinase 2 (LATS2) in non-small cell lung cancer cells by recruiting EZH2 to the LATS2 promoter [28]. We searched HiMoRNA for PVT1-correlated peaks around the LATS2 gene. We detected only H3K4me3 peaks that were negatively correlated with PVT1, which appears reasonable since PVT1 attracts EZH2 and participates in establishing the repressive H3K27me3 mark. In RNA-Chrom we observed contacts around one of these peaks (peak_169403, chr13:21045571-21046978) in two experiments for two cell lines (K562; MDA-MB-231). The Genome Browser visualization confirms the presence of this peak in the promoter of LATS2 (a step-by-step analysis is provided in Supplementary Table 3). An additional validation for PVT1 regulation of LATS2 comes from the EZH2-mediated Red-ChIP data (hES cell). A peak of PVT1 contacts (chr13:21168000-21224000, qValue=0.09) obtained by BaRDIC is located 97 Kb upstream of LATS2.

The lack of a positive correlation between lncRNA PVT1 and the ChIP-seq H3K27me3 peaks in the HiMoRNA seems to be due to overly strict peak selection. Still, this case is in concordance with previously reported results.

## 4. Discussion

HiMoRNA originally provides a collection of lncRNA–genomic loci for which lncRNA expression is significantly correlated with the histone modification signal across multiple cell and tissue types. Since HiMoRNA currently contains over five million correlations for ten histone modifications and 4145 lncRNA, it is expected that some of these correlations may be false positives; therefore, they might require additional experimental validation. RNA-Chrom is a manually curated database that contains genome-wide RNA–chromatin interactions data. Unfortunately, experimental data on RNA-chromatin interactions suffer from RNA expression bias, underrepresentation of contacts for low expressed lncRNAs, overrepresentation of contacts for the nascent transcripts, and do not provide mechanistic hypotheses. Having this in mind, it is reasonable to integrate HiMoRNA and RNA-Chrom databases to characterize the effects of lncRNA on epigenetic modifications and gene expression.

In order to ascertain the functional role of RNA in the corresponding DNA locus, additional genome-wide data and annotations are required. These may include information on the structure of chromatin, gene expression, or the localization of DNA-binding and chromatin-modifying proteins. RNA-Chrom provides a variety of information about the interaction of RNA with chromatin, which can be used in a comparative analysis with other data or as a target for experimental refinements.

As mentioned above, no H3K27ac or H3K4me1 peaks were observed in the FABP4 region when searching the HiMoRNA database. This finding contradicts experimental data and RNA-Chrom DB. There are other missing instances from both HiMoRNA and RNA-Chrom for known cases of lncRNA action in epigenetic regulation and chromatin. For example, no MEG3-correlated H3K27me3 peaks were observed in HiMoRNA, even though MEG3 has been reported as a regulator of PRC2 and is involved in H3K27me3 maintenance, particularly in the promoter regions of SMAD2, TGFB2, and TGFBR1 genes [8]. The absence of such peaks can be attributed to the fact that the majority of lncRNAs are expressed in a cell-specific manner, and the observed regulation was reported in a cell type that is not present in HiMoRNA. Even when ChIP-seq data are available, the standard peak calling procedure may be overly robust, potentially leading to the loss of biologically significant interactions.

We also encountered situations where a prediction from the HiMoRNA database was supported by the literature, but we did not detect RNA-chromatin contacts in the corresponding region in RNA-Chrom. For example, the majority of MAPKAPK5-AS1 peaks are not supported by MAPKAPK5-AS1 contacts with chromatin. We believe that such cases are due to the low number of detected contacts for a given RNA and the lack of data in a given cell type.

Since both databases have their own biases, it is possible that some known cases might be missing from the integration. Nevertheless, we posit that the integration of the databases reduces various biases owing to the multi-omics nature of the integrated data, thereby enabling the generation of interpretable hypotheses on the mechanisms of lncRNA epigenetic regulation of gene expression, supported by experimental data on RNA-chromatin interactions.

## 5. Future Development

The field of functional lncRNA annotation is a rapidly developing area of research. We are committed to maintaining the integration of HiMoRNA and RNA-Chrom as each database is updated with new data on lncRNA function in epigenetic regulation and interaction with chromatin. As both databases evolve and improve, their integration will also improve. As more experimentally validated data becomes available, we aim to build several predictive models for lncRNA histone modification. In addition, we will continue to improve the performance of our computer servers for storing and analyzing the newly generated data. It is expected that these ongoing efforts and our dedication to the development and enhancement of the integration of HiMoRNA and RNA-Chrom will facilitate a deeper understanding of lncRNA-mediated chromatin regulation.

## 6. Acknowledgments

The authors would like to thank Arina Nikolskaya for providing the Red-ChIP processed data, as well as the anonymous reviewers for their valuable suggestions.

## Author Contributions

Improved HiMoRNA’s interface and database, software development, I.I.; adaptation of the RNA-Chrom web resource for integration with HiMoRNA, gene names correspondence, project administration, G.R.; usecases, G.R. and D.M.; general supervision, A.M. and Y.M.; manuscript preparation, I.I., G.R., D.M., A.M. and Y.M. All authors have read and agreed to the published version of the manuscript.

## Disclaimer/Publisher’s Note

The statements, opinions and data contained in all publications are solely those of the individual author(s) and contributor(s) and not of MDPI and/or the editor(s). MDPI and/or the editor(s) disclaim responsibility for any injury to people or property resulting from any ideas, methods, instructions or products referred to in the content.

