## Supplementary Table 2 for "HiMoRNA and RNA-Chrom integration: Chromatin-Associated LncRNAs in Genome-Wide Epigenetic Regulation"

**Supplementary Table 2.** Step-by-step analysis of lncRNA MIR31HG using integrated HiMoRNA and RNA-Chrom databases

| **Web resource** | **Step** | **Current page** | **Actions** | **Resulting page** |
|---|---|---|---|---|
| HiMoRNA | 1 | HiMoRNA welcome page | click on the “Search page” button or click on the “Search” button on the toolbar | Search page |
| | 2 | Search page | add “MIR31HG” to the corresponding field<br>select histone modifications of interest (H3K4me1, H3K27ac)<br>add genomic coordinates for upstream-extended GLI2 (chr2 120725622 120992653)<br>click on the “Search” button | Search result page |
| | Optional | Search result page | click on the histone modification hyperlink | Histone modifications page |
| | | | click on the lncRNA hyperlink | lncRNA page |
| | | | click on the gene hyperlink | Modification-associated gene page |
| | | | click on the correlation coefficient hyperlink | Peak-lncRNA`s correlation page |
| | | | click on the “Download” button | Download the results table |
| | 3 | Results table page | • choose the peak of interest with the radio button on the left side<br><br>click on the “Go to RNA-Chrom DB” button, then on the “MIR31HG RNA contacts with the locus (peak_473655)” button and select the peak extension | Graphic summary page containing contacts of lncRNA MIR31HG with chromatin in the peak region |
| RNA-Chrom | 4 | Graphic summary page | choose the particular experiment with checkboxes<br>click on the “ALL TARGET GENES” button | MIR31HG RNA target genes (in the locus of the extended peak) page |
