## Supplementary Table 3 for "HiMoRNA and RNA-Chrom integration: Chromatin-Associated LncRNAs in Genome-Wide Epigenetic Regulation"

**Supplementary Table 3.** Step-by-step analysis of lncRNA PVT1 using integrated HiMoRNA and RNA-Chrom databases

| Web resource | Step | Current page | Actions | Resulting page |
|---|---|---|---|---|
| HiMoRNA | 1 | HiMoRNA welcome page | - click on the “Search page” button or click on the “Search” button on the toolbar | Search page |
| | 2 | Search page | - add “PVT1” to the corresponding field<br>- select all histone modifications<br>- add "LATS2" to the "Gene/Gene ID" field<br>- click on the “Search” button | Search result page |
| | 3 | Results table page | - choose the peak of interest with the radio button on the left side (e.g. peak_169403)<br>- click on the “Go to RNA-Chrom DB” button, then on the “PVT1 RNA contacts with the locus (peak_169403)” button and select the peak extension (e.g. 25000) | Graphic summary page containing contacts of lncRNA PVT1 with chromatin in the peak region |
| RNA-Chrom | 4 | Graphic summary page | - choose the particular experiment with checkboxes (Exp.IDs: 8, 10)<br>- click on the “VIEW IN GENOME BROWSER” button | UCSC Genome Browser |
