## Supplementary for "HiMoRNA and RNA-Chrom integration: Chromatin-Associated LncRNAs in Genome-Wide Epigenetic Regulation"

**Supplementary Table 1.** LncRNA correspondence table.

**Supplementary Table 2.** Step-by-step analysis of lncRNA MIR31HG using integrated HiMoRNA and RNA-Chrom databases.

**Supplementary Table 3.** Step-by-step analysis of lncRNA PVT1 using integrated HiMoRNA and RNA-Chrom databases.

**Supplementary Figure 1.** Intersection of 4145 genes from HiMoRNA DB with 60619 genes from RNA-Chrom DB. A. Dividing gene pairs into 6 groups depending on the similarity metrics they satisfy. Those groups in which one-to-one correspondence between genes is achieved are highlighted in red. B. Number of gene pairs in groups 2, 4, 5 and 6.

**Supplementary Figure 2.** Illustration of HiMoRNA and RNA-Chrom DB usage on the example of MIR31HG usecase. A. Creation of a query for MIR31HG and its target histone marks in HiMoRNA. B. HiMoRNA`s table with search results. C. Switching to RNA-Chrom DB. D. RNA-Chrom DB page with MIR31HG-chromatin contacts in the extended peak locus. E. RNA-Chrom DB`s table with all target genes for Exp.ID 9.
