## Supplementary figures and images for "HiMoRNA and RNA-Chrom integration: Chromatin-Associated LncRNAs in Genome-Wide Epigenetic Regulation"

### Supplemental Figure 1

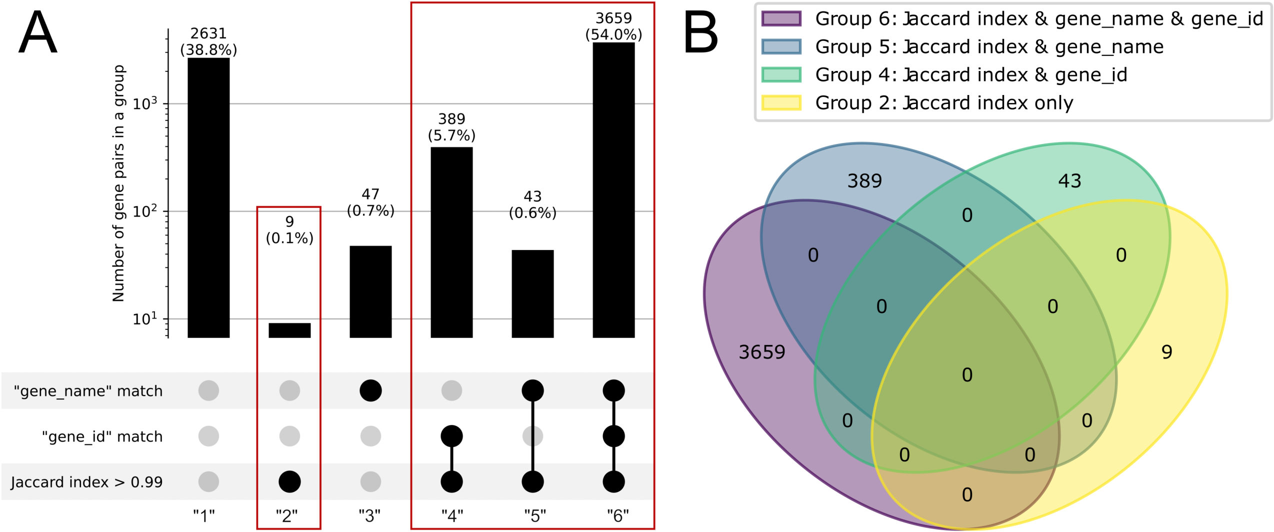

### Supplemental Figure 2

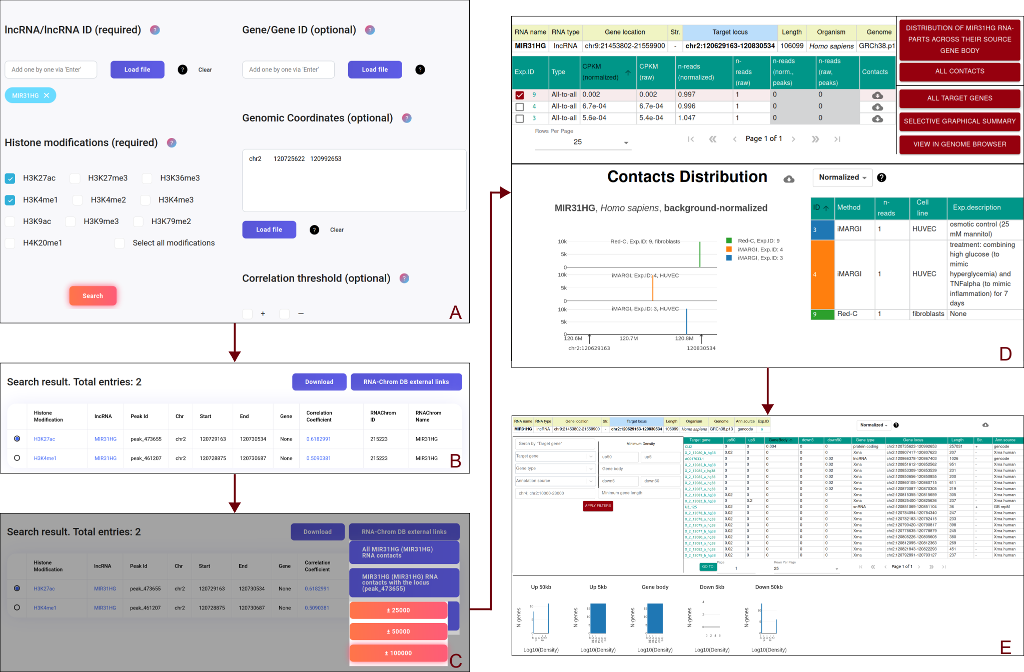
